# A roadmap for the characterization of energy metabolism in human cardiomyocytes derived from induced pluripotent stem cells

**DOI:** 10.1101/2020.09.28.316745

**Authors:** Giulia Emanuelli, Anna Zoccarato, Angelos Papadopoulos, Mei Chong, Matteo Beretta, Kai Betteridge, Katrin Streckfuss-Bömeke, Ajay M Shah

## Abstract

Human induced pluripotent stem cell-derived cardiomyocytes (hiPSC-CM) are an increasingly employed model in cardiac research and drug discovery. A significant limitation with respect to clinical translation is their immature structural and functional phenotype. Cellular metabolism plays an integral role in determining phenotype but the metabolic profile of hiPSC-CM during maturation is poorly characterized. In this study we employ a combination of methods including extracellular flux and ^13^C-glucose flux analyses to study the metabolic profile of hiPSC-CM over a 12 week maturation period. Results show a progressive remodeling of pathways involved in energy metabolism and substrate utilization. The oxidative capacity of hiPSC-CM and particularly their ability to utilize fatty acids increased with time. In parallel, relative glucose oxidation was reduced while glutamine oxidation was maintained at similar levels. Anaerobic glycolysis as assessed by lactate production was maintained throughout the 12 week period but with significant alterations in proximal glycolytic enzymes such as hexokinase and phosphofructokinase. We also observed a progressive maturation of mitochondrial oxidative capacity at comparable levels of mitochondrial content between timepoints. The time-dependent restructuring of the hiPSC-CM metabolic profile indicates their potential to overcome the lack of full maturation previously reported and enhance their applicability for pharmacological studies and cardiac disease modeling.

## Introduction

The ability to derive cardiomyocytes (CM) from induced pluripotent stem cells (iPSC) has opened new avenues for drug discovery and disease modeling using human cells. hiPSC-derived CM (hiPSC-CM) exhibit many features of primary cardiomyocyte structure, electrophysiology, contraction and pharmacological responses to inotropic agents (Otsuji et al., 2010; Paik et al., 2020; Robertson et al., 2013). However, these cells in general display an overall immature phenotype, with their transcriptome more closely resembling fetal than adult cardiomyocytes (van den Berg et al., 2015). hiPSC-CM retain a higher proliferative potential than adult CM, a less organized sarcomere structure, spontaneous contractile activity and slower action potentials (Ivashchenko et al., 2013). This immature phenotype constitutes the main limitation to the use of these cells in disease modeling and potentially regenerative medicine. As such, considerable efforts have been made to develop strategies to induce faster or more complete maturation, including electrical and mechanical activation and other measures (Kolanowski et al., 2017; Machiraju and Greenway, 2019). Nevertheless, the molecular mechanisms determining the lack of full maturation of hiPSC-CM are poorly understood.

In contrast to the transcriptomic, electrophysiological and contractile features of hiPSC-CM, there has been limited research around their metabolic profile and the changes accompanying maturation. This may be a critical aspect because metabolic processes are inextricably linked to structural and functional maturation of all cells. In mammalian CM, it is well recognized that pathways involved in energy metabolism undergo profound remodeling during the perinatal period (Lopaschuk and Jaswal, 2010). Anaerobic glycolysis supports the anabolic requirements of the proliferative state during fetal life (Hu et al., 2016) as well as providing ATP in the relatively hypoxic environment at that stage (Ezashi et al., 2005). After birth, there is a switch to oxidative energy production which is thought to also be important for the transition of proliferating immature CM to terminally differentiated cells (Puente et al., 2014). The terminal differentiation of CM is accompanied by mitochondrial maturation, with an increase in content, development of a tubular structure with elongated cristae and enhanced polarization (Cho et al., 2006). The substrates that are used by CM for energy production vary substantially depending upon the maturation stage. In the fetus, the majority of oxygen consumption is attributable to oxidation of lactate, which is abundantly provided by the placental tissue (Burd et al., 1975). After birth, with increased oxygen availability, there is a major shift from anaerobic glycolysis to fatty acid (FA) oxidation (Lopaschuk et al., 1991). Glucose oxidation is estimated to contribute 15-20% of energy by the stage of weaning, similar to the metabolic profile of the adult heart (Lopaschuk and Jaswal, 2010).

Previous studies have reported that hiPSC-CM have an immature fetal-like metabolic phenotype, even after 1 month after *in vitro* differentiation (Robertson et al., 2013), associated with high rates of glycolysis and very low oxidation potential (Rana et al., 2012). However, changes during maturation have not been characterized in detail. Energy metabolism is also tightly intertwined with adult cardiomyocyte remodeling in pathological conditions such as chronic ischemia and chronic pressure or volume overload (Doenst et al., 2013). It is therefore important to have a well-defined baseline metabolic profile if hiPSC-CM are to be used to model such pathologies. In this work, we have undertaken a detailed analysis of the metabolic remodeling that occurs in hiPSC-CM with their prolonged culture. We studied the metabolic profile of hiPSC-CM from healthy donors at an early stage (6 weeks) commonly employed by many investigators, and a late stage (12 weeks) after CM generation. We used a variety of different readouts to assess anaerobic glycolysis, mitochondrial respiration, polarization, structure, metabolic flux, and gene expression and protein levels of key enzymes. We report a roadmap of the metabolic changes that occur in hiPSC-CM during maturation after their initial derivation from iPSC.

## Results

### Prolonged culture aids structural maturation of hiPSC-CM

The hiPSC cell lines FB2 and Ctrl2 (Borchert et al., 2017; Wolf et al., 2013) both demonstrated features of stemness and pluripotency as assessed by typical morphology, alkaline phosphatase (ALP) activity and the expression of the selfrenewal markers NANOG, SOX2, LIN28, TRA-1-60 and SSEA-4 assessed by immunofluorescence (Supplementary Fig. 1A-B). The differentiation into hiPSC-CM (Lian et al., 2012) was accompanied by the repression of pluripotency and selfrenewal genes, followed by an early transient increase of stage-specific early cardiogenic factors (i.e. *MIXL1 and GATA4*) and a significant increase of the cardiac-specific markers *ACTN2* and *MY2* (Supplementary Fig. 1C-F). At week 6 (termed early-stage hiPSC-CM), the CM purity as assessed by α-actinin positive (α-Actinin^+^) cells by FACS was >95% (Fig. 1A). With continued culture to 12 weeks (termed late-stage hiPSC-CM), there was a significant increase in the proportion of cells positive for cardiac troponin T (cTnT, from 49.2 to 89.2 %) and myosin light chain-2a (MLC2a, from 7.8 to 21.2 %). There was also a substantial increase in the intensity of α-Actinin, cTnT and MLC2v fluorescence, indicative of ongoing CM maturation throughout this period, whereas MLC2a was not upregulated during maturation (Fig. 1A). These results are consistent with previous reports (Bedada et al., 2016; Kamakura et al., 2013). Sarcomere structure as assessed by co-staining cells with antibodies against C-terminal or N-terminal portions of titin and α-Actinin was similar in early-stage and late-stage hiPSC-CM (Fig. 1B), with a consistent pattern of localization of these myofilament proteins (Fig. 1C). The sarcomere length and regularity were also similar between these stages (Fig. 1D).

**Figure 1.**
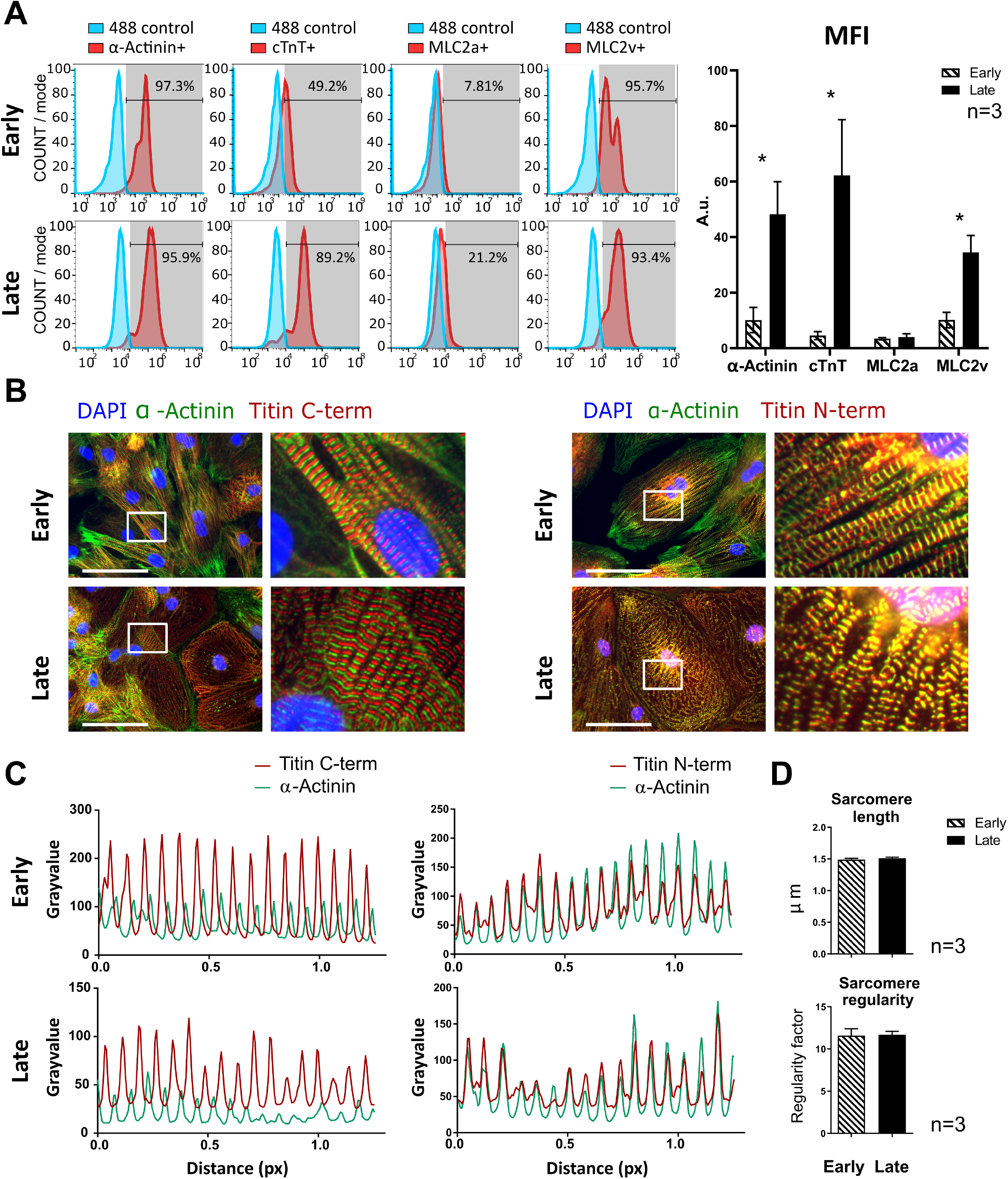
Effect of prolonged culture of hiPSC-CM on sarcomeric protein levels and ultrastructure. A) FACS analysis of hiPSC-CM at early-stage (week 6) and late-stage (week 12). Left panel: positive cells in the cell population (%); right panel: mean fluorescence intensity (MFI) ± S.E.M. *, p<0.05 by unpaired t-tests; n=3 differentiation experiments. B) Representative immunofluorescence images; scale bar, 200 μm. C) Co-localisation patterns of immunofluorescence for the markers shown. The Pearson correlation coefficients for the data shown were: Titin C-term (C-terminal) and α-Actinin, early r=-0.360, late r=-0.448; Titin N-term and α-Actinin, early r=0.861, late r=0.853. number of pairs=189. D) Quantification of sarcomeric parameters from the immunofluorescence images represented in B (α-Actinin signals), processed via Fast Fourier Transformation. n=3 independent experiments, 10 cells/experiment.

### Restructuring of the glycolytic network

Fetal CM are thought to be highly reliant on anaerobic glycolysis for energy production but enzymatic and topological restructuring of the glycolytic network is reported to be necessary for the differentiation and maturation of CM towards oxidative energy metabolism (Chung et al., 2010). We compared features of glycolysis and related pathways between early-stage and late-stage hiPSC-CM. The quantification of extracellular acidification rate (ECAR) on a Seahorse flux analyzer as an index of lactate production showed no differences between early-stage and late-stage hiPSC-CM (Fig. 2A). There was also no difference in the glycolytic reserve between stages, as assessed by a glycolytic stress test in which complex V (ATP synthase) is inhibited with oligomycin (Fig. 2A).

**Figure 2.**
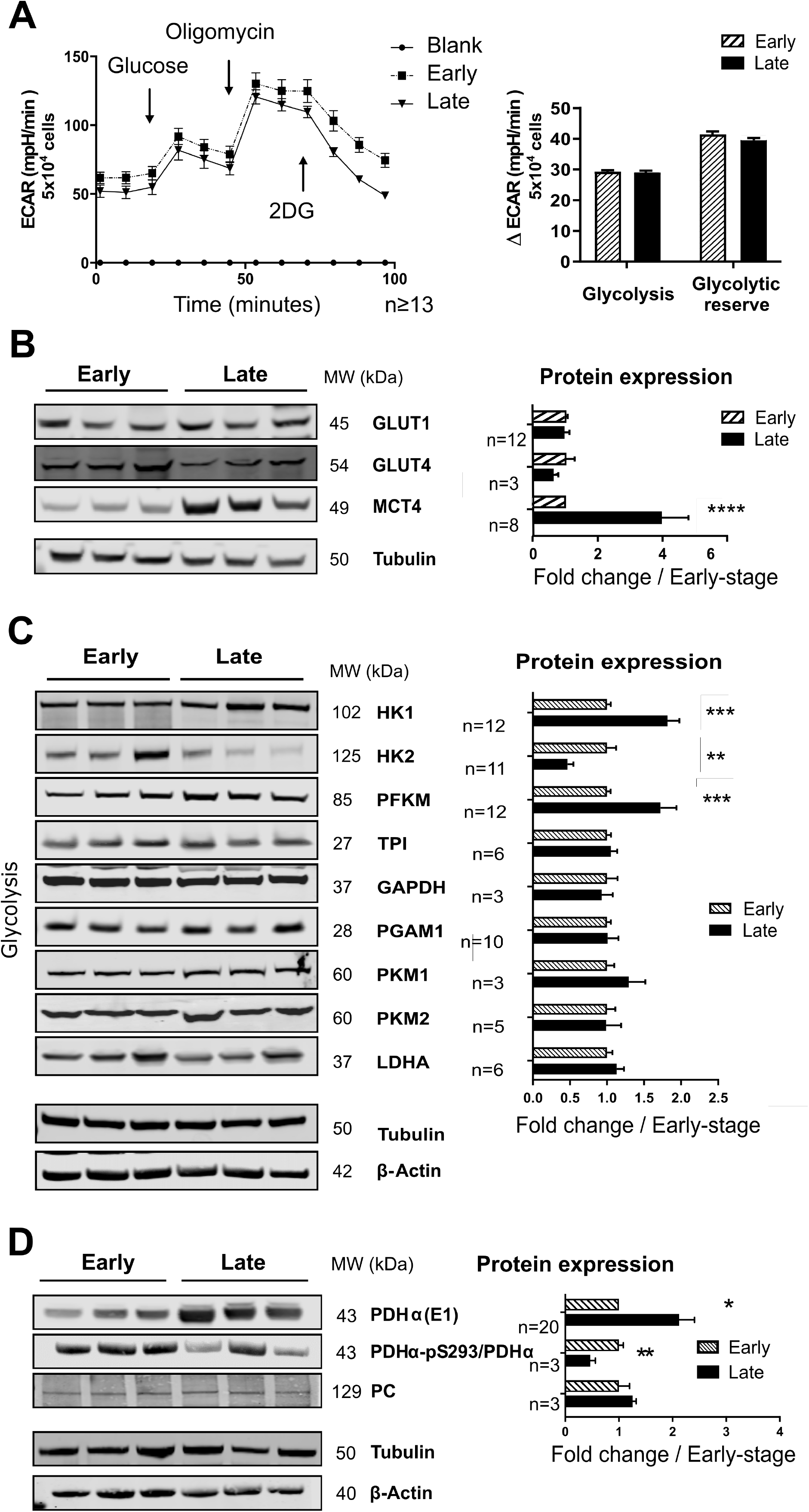
The glycolytic network in maturing hiPSC-CM. A) Glycolysis stress tests in early-stage versus late-stage hiPSC-CM, performed on a Seahorse analyzer. Left panel: Average profiles of the extracellular acidification rate (ECAR). 2DG, 2-deoxyglucose. Right panel: Quantification of glycolytic parameters. B) Left panel: Representative western blots for carbohydrate plasma membrane facilitated transporters. Right panel: Quantification of relative protein expression. C) Left panel: Representative western blots for glycolytic enzymes. Right panel: Quantification of relative protein expression. D) Left panel: Representative western blots for pyruvate converting enzymes. Right panel: Quantification of relative protein expression. All data are mean ± S.E.M. **, p<0.01; ***, p<0.001; ****, p<0.0001 by unpaired t-tests. Numbers as indicated in figure (3-20). SLC: solute carrier (family); HK: hexokinase; PFKM: muscle phosphofructokinase; TPI: triosephosphate isomerase; GAPDH: glyceraldehyde-3-phosphate dehydrogenase; PGAM: 2,3-bisphosphoglycerate-independent phosphoglycerate mutase; PKM: pyruvate kinase M; LDHA: lactate dehydrogenase subunit A; PDH: pyruvate dehydrogenase; PC: pyruvate carboxylase.

We further characterized glucose utilization. The protein levels of the glucose transporters, GLUT1 (SLC2A1) and GLUT4 (SLC2A4), were similar in early-stage and late-stage cells. However, levels of the lactate transporter MCT4 (SLC16A3) were 4-fold higher in late-stage compared to early-stage hiPSC-CM (Fig. 2B). We assessed the protein levels of a panel of enzymes involved in glycolysis (Fig. 2C). This showed a significant isoform shift from hexokinase 2 (HK2) to hexokinase 1 (HK1), and an increase in muscle phosphofructokinase (PFKM), in late-stage cells. No significant changes were observed in the other main glycolytic pathway enzymes. Hexokinase and PFKM are proximal rate-limiting enzymes in the glycolytic pathway which may influence the partitioning of glycolytic intermediates between forward glycolysis and entry into the tricarboxylic acid (TCA) cycle or other glycolytic branch pathways.

Glycolytic flux assessed by the ^13^C enrichment of [1,2,3]-^13^C lactate in cells incubated with U-^13^C glucose was >97% at both early and late stages, as well as undifferentiated hiPSCs, indicating similarly high level of glycolytic flux (Fig. 3A). To assess flux into the pentose phosphate pathway (PPP), we quantified the [2,3]^13^C lactate isotopomer in cells incubated with U-^13^C glucose. This analysis relies on the re-entry of 3-phosphogycerate from the PPP into glycolysis and the subsequent generation of lactate (Goodwin et al., 2001) (Fig. 3A). The estimated PPP flux was found to be very low both in early-stage and late-stage hiPSC-CM and not significantly different from undifferentiated iPSC. The protein levels of the two major PPP regulatory enzymes, glucose 6-phosphate dehydrogenase (G6PD) and 6-phosphogluconate dehydrogenase (PGD), were significantly lower in late-stage compared to early-stage hiPSC-CM (Fig. 3B). The hexosamine biosynthetic pathway (HBP), another glycolytic branch pathway, results in the formation of uridine diphosphate-N-acetylglucosamine (UDP-GlcNAc) which is used in post-translational O-GlcNAcylation of proteins (Ngoh et al., 2010). The level of overall protein O-GlcNAcylation was ~1.7-fold higher in late-stage compared to early-stage hiPSC-CM (Fig. 3C), suggesting increased HBP activity. This was associated with a relative change in isoforms of the rate-limiting HBP enzyme, glutamine-fructose-6-phosphate amidotransferase (GFAT), from isoform 2 to 1, with a significant reduction in GFAT2 in late-stage hiPSC-CM (Fig. 3B).

**Figure 3.**
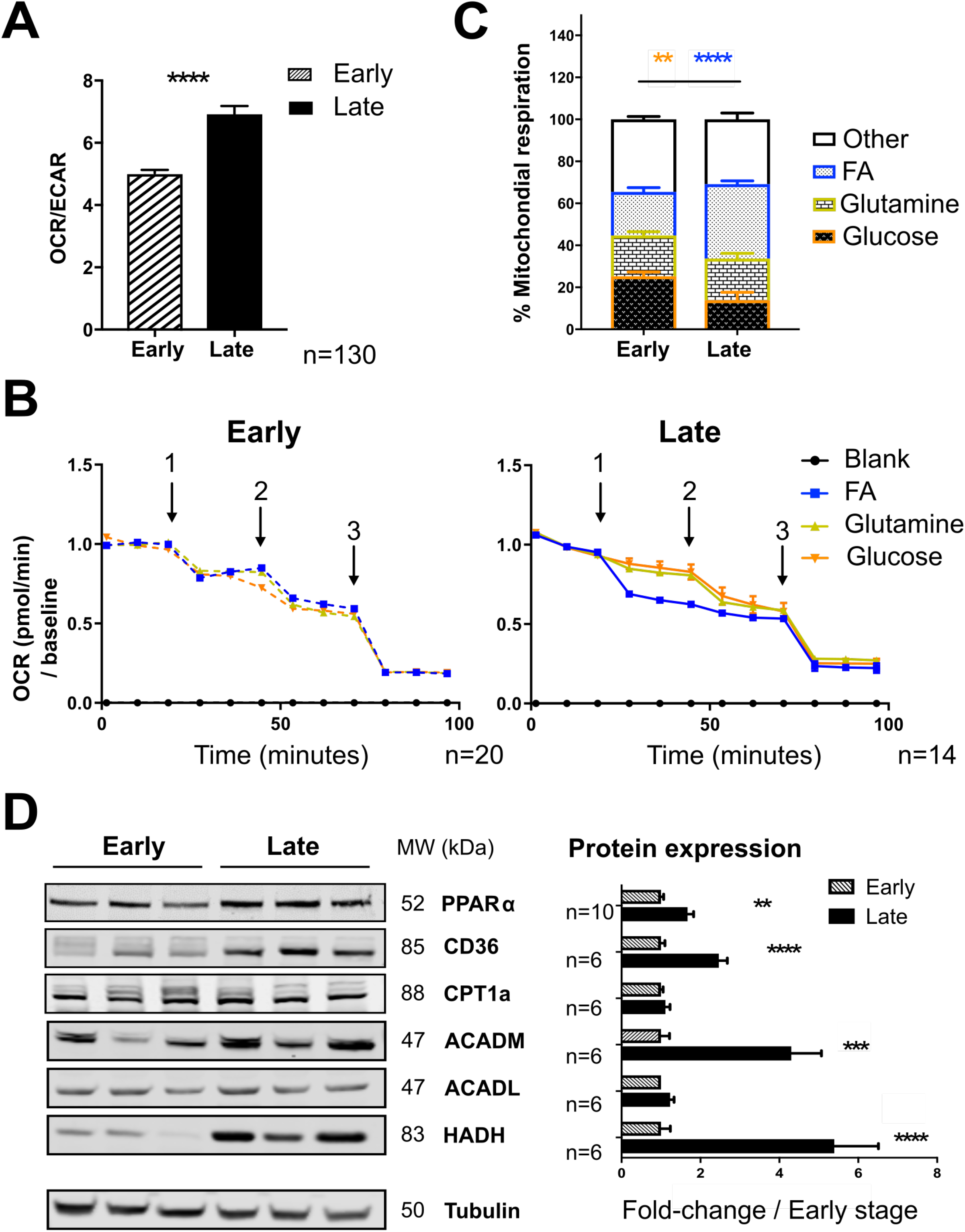
Characterization of glycolytic branch pathways. A) Left: Schematic representation of lactate labeling patterns after metabolism of U-^13^C glucose through the glycolytic or pentose phosphate pathway (PPP). Right: Quantification of the % ^13^C labeling of [1,2,3]^13^C-lactate and [2,3]^13^C-lactate to estimate glycolysis flux and PPP flux respectively. Cells were incubated in 10 mM U-^13^C Glucose for 16 h. Metabolite quantification was undertaken by HSQC NMR spectroscopy. Data are mean ± S.E.M; n=3. B-C) Left panels: Representative western blots for the targets shown. Right panels: Quantification of relative protein expression; data are mean ± S.E.M. *, p<0.05; **, p<0.01 by unpaired t-tests. n=3-7 as indicated in figure. PPP: pentose phosphate pathway; GFAT: glutamine - fructose-6-phosphate transaminase; OGT: O-linked N-acetylglucosamine (GlcNAc) transferase; G6PD: glucose-6-phosphate dehydrogenase; PGD: phosphogluconate dehydrogenase.

Turning to the entry of glycolytic intermediates into the TCA cycle, the expression of the pyruvate dehydrogenase alpha subunit (PDHαE1) was 2-fold higher in late-stage hiPSC-CM whereas inhibitory phosphorylation at S293 (site1) (Rardin et al., 2009) was substantially lower (Fig. 2D). We also found a marked increase in pyruvate dehydrogenase kinase-4 (PDK4) in late-stage hiPSC-CM and a smaller increase in pyruvate dehydrogenase phosphatase-1 (PDP1) but no change in PDK1 levels (Supplementary Fig. 2A). Protein levels of pyruvate carboxylase (PC), which may support anaplerotic entry of intermediates into the TCA cycle, were similar in early-stage and late-stage cells (Fig. 2D).

### Oxidative metabolism and substrate preference

To assess the relative balance between anaerobic glycolysis and oxidative metabolism, hiPSC-CM were incubated in RPMI 1640 medium supplemented with glucose, FA, L-carnitine and glutamine. The oxygen consumption rate (OCR) and ECAR were quantified simultaneously on a Seahorse instrument. This revealed that the OCR/ECAR ratio was significantly higher in late-stage compared to early-stage cells (Fig. 4A), indicating a time-dependent shift towards a more oxidative metabolic profile.

**Figure 4.**
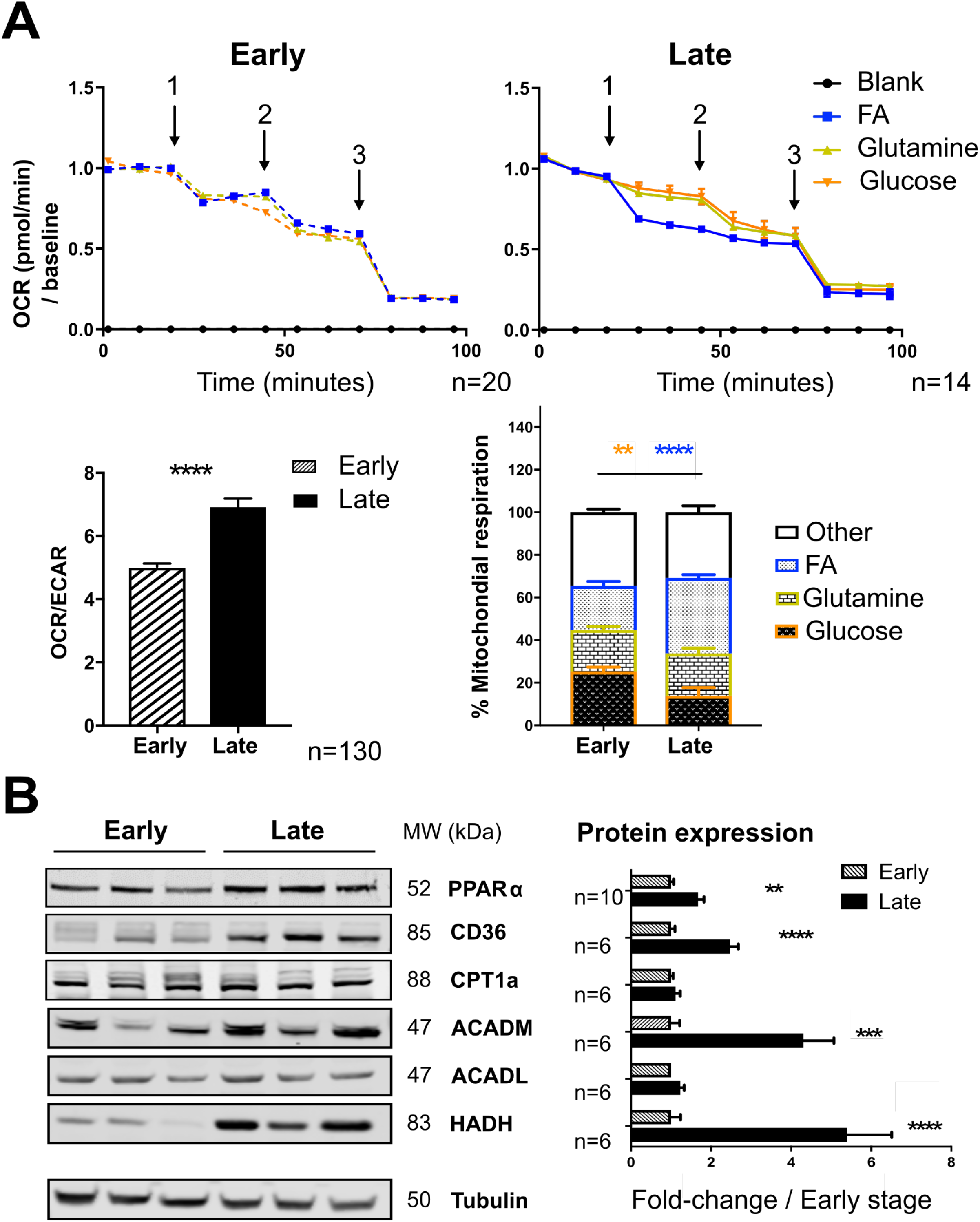
Relative anaerobic versus oxidative metabolism and alterations in substrate utilization in hiPSC-CM. A) Ratio of oxygen consumption rate (OCR) over extracellular acidification rate (ECAR) in early-stage versus late stage hiPSC-CM equilibrated in complete medium. Data are mean ± S.E.M. ****, p<0.0001 by Student’s t-test. n=130 (cumulative readings from all Fuel Source tests performed). B) In Fuel Source tests, the utilization of each substrate was targeted by a pharmacological inhibitor. The substrates were fatty acids (FA), glutamine and glucose. A single inhibitor was added at time-point 1; the other two inhibitors were added at time-point 2 (all combinations were performed in parallel on the same plate); and Antimycin A and Rotenone were added at time-point 3 (assay summary in Supp. Table 4). The graphs show the average profiles of OCR. C) Quantification of the contribution of different substrates to total OCR. Data are mean ± S.E.M. **, p<0.01 and ****, p<0.0001 by 2 way ANOVA followed by Tukey’s test. n=20 (early); n=14 (late). The colors of the asterisks indicate the substrate being considered for early vs late comparison. C) Left panel: Representative western blots for targets involved in FA metabolism. Right panel: Quantification of relative protein expression. Data are mean ± S.E.M. **, p<0.01; ***, p<0.001; ****, p<0.0001 by unpaired t-tests. n=6-10 as indicated in figure. PPARα: peroxisome proliferator activated receptor alpha; CD36: cluster of differentiation, molecule 36; CPT1a: carnitine-palmitoyl transferase subunit 1a; ACADM/L: medium/light chain acyl-CoA dehydrogenase; HADH: hydroxyacyl-CoA dehydrogenase.

Next, we evaluated the relative contribution of different substrates to total oxidative metabolism. In cells incubated in complete medium (as above), OCR was measured in the presence of specific inhibitors that each targeted a single substrate. We used etomoxir to inhibit FA oxidation, BPTES to inhibit glutamine oxidation and UK5099 to inhibit the mitochondrial pyruvate carrier and therefore glucose oxidation, and then quantified the corresponding reductions in OCR (Fig. 4B). A single inhibitor was added in step 1, then the other two inhibitors in step 2 (each combination was performed in parallel on the same plate), and finally Antimycin A and rotenone in step 3 to completely inhibit the electron transport chain (ETC). A different substrate inhibitor was added in different wells but with all the assays performed on a single plate. Using this approach, we calculated the proportion of total OCR that was accounted for by each substrate in the complete medium. This analysis revealed that the relative contribution of FA oxidation increased significantly in late-stage compared to early-stage hiPSC-CM, at the expense of glucose oxidation (Fig. 4C). A significant amount of glutamine oxidation contributed to the overall OCR both in early-stage and late-stage cells.

The relative increase in FA oxidation was accompanied by an increase in levels of PPARα, the master transcription factor that drives lipid metabolism (Gilde, et al., 2003), in late-stage compared to early-stage cells (Fig. 4D). We also observed increased levels of the plasma membrane FA transporter CD36 and the mitochondrial β-oxidation enzymes, medium-chain acyl-CoA dehydrogenase (ACADM) and 3-hydroxyacyl-CoA dehydrogenase (HADH), in late-stage cells (Fig. 4D). The levels of carnitine-palmitoyl transferase 1 (CPT1a) were similar in early-stage and late-stage cells. Since CPT1 is sensitive to inhibition by malonyl-CoA, we quantified the levels of acetyl-CoA carboxylase (ACC1 and 2) and malonyl-CoA decarboxylase, which regulate malonyl-CoA levels (Foster, 2012). ACC2 increased in late-stage compared to early-stage hiPSC-CM but there were no differences in ACC1 or malonyl-CoA decarboxylase (Supplementary Fig. 2B). However, a slight shift in ACC1 electrophoretic mobility in late-stage hiPSC-CM samples might indicate differences in post-translational modification.

### Mitochondrial content, structure and function

Changes in mitochondrial structure and function are key drivers of metabolic maturation in the developing heart. We therefore compared mitochondrial parameters in early-stage and late-stage hiPSC-CM.

Ultrastructural examination of hiPSC-CM by transmission electron microscopy revealed differences in interfibrillar mitochondria between early-stage and late-stage cells (Fig. 5A). Cells at both stages had more elongated mitochondria and better-defined cristae than undifferentiated hiPSCs; however, late-stage hiPSC-CM had longer mitochondria and a more developed branching pattern. These changes are suggestive of a maturation of the mitochondrial network. To assess this at a functional level, we quantified the mitochondrial membrane potential (ΔΨ_mt_). This was found to be significantly higher in late-stage compared to early-stage cells (Fig. 5B). In addition, the basal mitochondrial calcium levels quantified using a mitochondrial-targeted FRET sensor were significantly higher in late-stage compared to early-stage hiPSC-CM (Fig. 5C). These results suggest the potential for higher activity of mitochondrial calcium-dependent dehydrogenases (such as PDH or TCA cycle enzymes) and a higher protonmotive force for oxidative ATP production.

**Figure 5.**
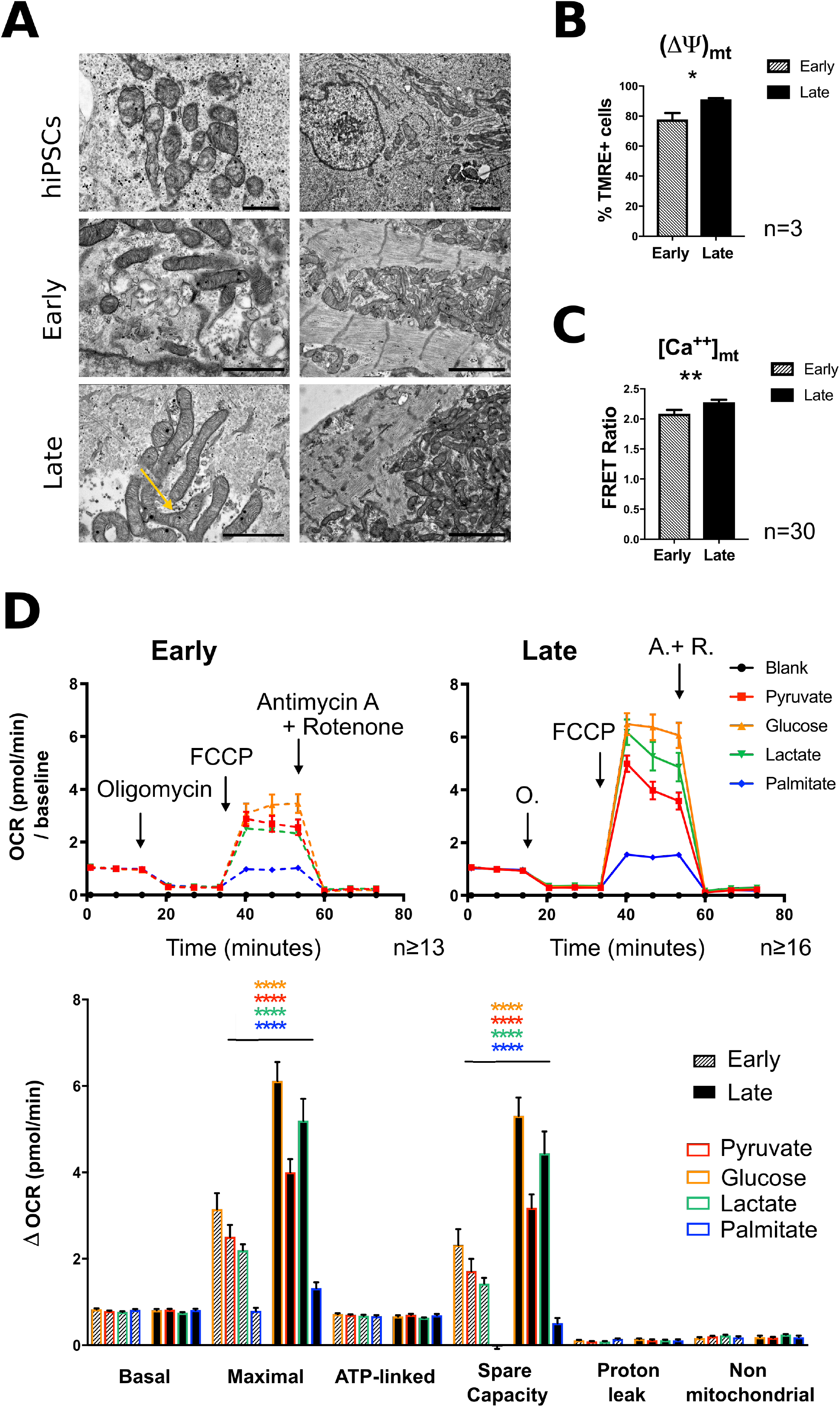
Mitochondrial ultrastructure and function in late-stage versus early-stage hiPSC-CM. A) Representative transmission electron micrographs of undifferentiated hiPSC and early-stage and late-stage hiPSC-CM showing interfibrillar mitochondria. Scale bars: left, 1 μm; right, 2 μm. Yellow arrow shows branching mitochondria. B) Quantification of (ΔΨ)_mt_ in TMRE-stained cells by flow cytometry. Data are mean ± S.E.M. *, p<0.05 by unpaired t-test. n=3. C) Quantification of [Ca^++^]_mt_ by a mitochondrial-targeted Cameleon FRET probe. Data are mean ± S.E.M. **, p<0.01 by unpaired t-test. n=30 (cumulative number of cells from 3 experiments). D) Mitochondrial stress tests in early-stage versus late-stage hiPSC-CM provided with a single substrate. Top panels show average OCR profiles. Bottom panels show quantification of parameters. Data are mean ± S.E.M. ****, p<0.0001 by 2-way ANOVA followed by Sidak’s multiple comparisons test. n ⩾ 13 (early); n ⩾ 16 (late). The colors of the asterisks indicate the substrate being considered for early vs late comparison.

To assess mitochondrial oxidative function, we measured the basal and maximal mitochondrial respiration of individual substrates (pyruvate, glucose, lactate or palmitate) in early-stage versus late-stage hiPSC-CM, using mitochondrial stress tests (Fig. 5D). These experiments revealed that the spare respiratory capacity (an index of the maximal respiration that can be achieved) was significantly higher in late-stage versus early-stage cells for each of the carbon sources tested. There were no significant differences between early-stage and late-stage cells in the basal OCR for each substrate nor in proton leak or non-mitochondrial oxygen consumption. These results are consistent with the features of mitochondrial maturation described above.

To assess whether these improvements in the mitochondrial oxidative capacity were due to altered mitochondrial content, we studied live cells labeled with Mitotracker® Green. We observed similar absolute mitochondrial area per cell or the fraction of the cell occupied by mitochondria (Fig. 6A). The ratio between nuclear and mitochondrial DNA was also similar between time-points (Fig. 6B). The protein levels and phosphorylation of the transcription factor peroxisome proliferator-activated receptor gamma coactivator 1-alpha (PGC1α), involved in mitochondrial biogenesis (Lehman et al., 2000), were no different in late-stage compared to early-stage cells (Fig. 6C). Finally, the protein levels of different components of the ETC were also similar in late-stage and early-stage cells (Fig. 6D). Collectively, these results indicate that mitochondrial content remains stable during the maturation of hiPSC-CM from 6 weeks to 12 weeks post-differentiation.

**Figure 6.**
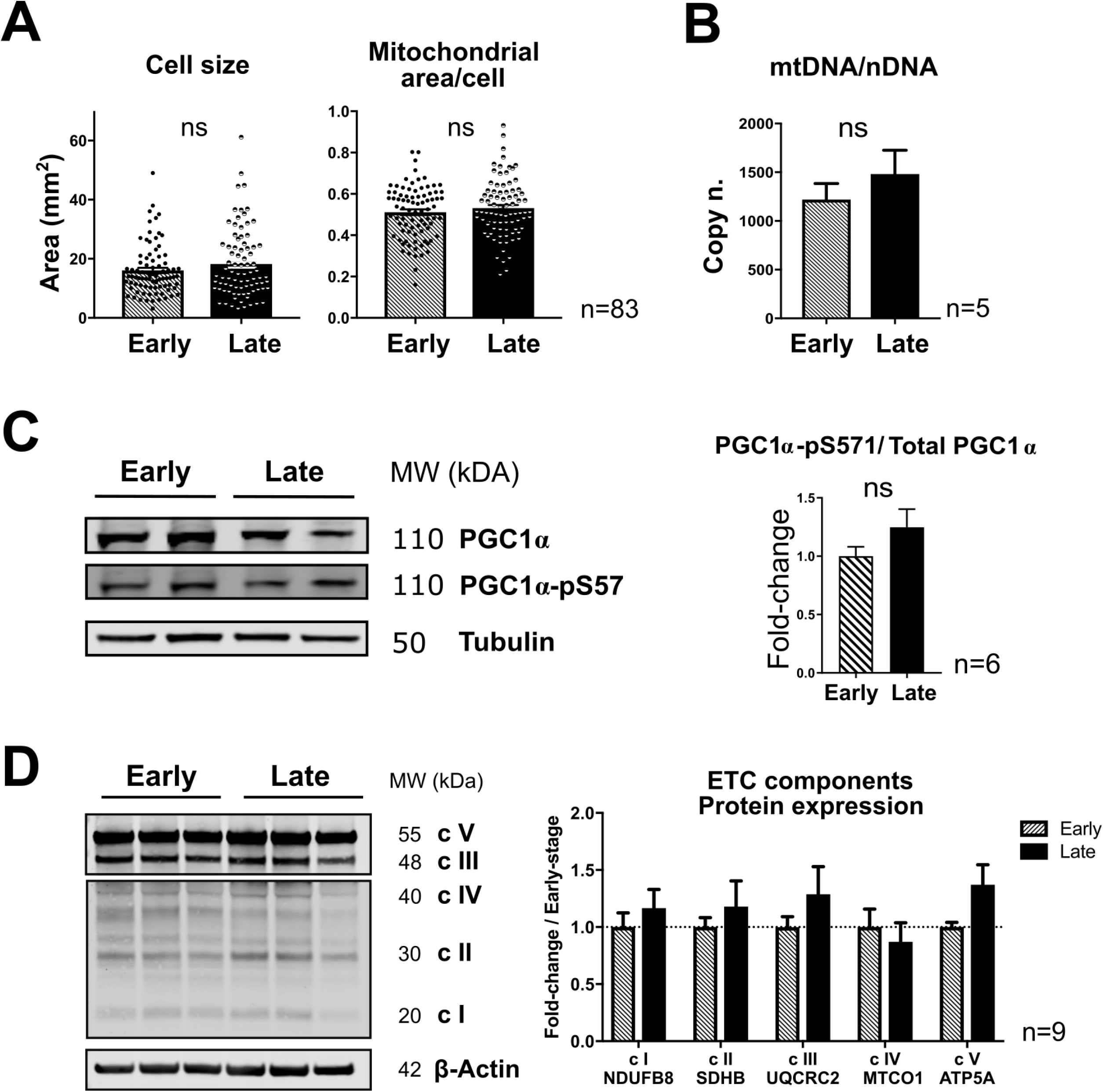
Mitochondrial content in early-stage and late-stage hiPSC-CM. A) Quantification of absolute cell area (left) or the fraction of mitochondrial area/cell (right) in cells loaded with Mitotracker® Green. Ns, p= 0.162; p = 0.336; not significant by unpaired t-test. n=83 (cumulative cells from 3 experiments). B) Mitochondrial to nuclear DNA ratio. Data are mean ± S.E.M. not significant by paired t-test; p=0.386; n=5 differentiation experiments. C-D). Left panels show representative western blots for the indicated proteins. Right panels show quantification of blots. Data are mean ± S.E.M. n=6 (C); n=9 (D). PGC1α: PPARG (peroxisome proliferator activated receptor gamma) coactivator 1 alpha; ETC: electron transport chain; NDUFB8: NADH:ubiquinone oxidoreductase subunit B8; SDHB, succinate dehydrogenase subunit B; UQCRC2: ubiquinol cytochrome c reductase core protein 2; MTCO1: mitochondrially encoded cytochrome c oxidase I; ATP5A: ATP synthase subunit alpha.

## Discussion

This study constitutes an effort towards a rigorous characterization of the metabolic profile of hiPSC-CM during their maturation after differentiation from iPSC. The remarkable potential of this cell model for drug discovery, disease modeling and regenerative medicine resides in its person-specific origin. A major limitation of the model, however, is the relatively immature state of hiPSC-CM as compared to fully differentiated adult CM. Energy metabolism is of fundamental importance in maintaining CM and heart function and is also integrally linked to physiological and pathophysiological CM growth and remodeling. As such, the metabolic profile of hiPSC-CM is a key factor to consider in the use of this cell model.

We identified significant changes in metabolic phenotype between early-stage and late-stage hiPSC-CM resembling some of the changes that are known to occur during developmental maturation of the mammalian heart. Anaerobic glycolysis as assessed either by U^13^C-labeled glucose experiments or quantification of ECAR using a Seahorse extracellular flux analyzer revealed similar rates in early-stage and late-stage hiPSC-CM. However, this was paralleled by an overall increase in oxidative metabolism such that the ratio between OCR and ECAR was higher in late-stage myocytes. Anaerobic metabolism is prevalent in stem cells and is associated with the proliferative state (Hu et al., 2016). The sustained rate of anaerobic glycolysis in hiPSC-CM, comparable to undifferentiated hiPSC, may be related to the anabolic requirements of these cells and could be involved in maintaining an immature phenotype. This hypothesis is supported by recent data showing that inhibition of HIF1-α or LDHA, which promote anaerobic metabolism, enhances the structural and functional maturation of hiPSC-CM (Hu et al., 2018). In the current study, it was of interest that although the overall forward glycolysis rates appeared similar between early-stage and late-stage hiPSC-CM, there were nevertheless significant changes in some of the key enzymes in the pathway. A switch in the expression from hexokinase 2 to hexokinase 1 occurred between early and late-stage hiPSC-CM. Hexokinase 1 associates with the mitochondrial fraction and is important in the neonatal period for the coupling of glycolysis to oxidative metabolism (Fritz et al., 1999) whereas hexokinase 2 is highly expressed in proliferating cells (such as stem cells and cancer cells). This was accompanied by evidence of altered functional activity of the glycolytic branch pathway, the HBP, during the maturation of hiPSC-CM – suggesting that more complex changes are occurring in the overall glycolytic network.

The utilization of different carbon substrates for oxidative phosphorylation and energy production, and the changes in relative substrate usage after birth, are key features of heart metabolism. We developed a novel protocol to test the relative contribution of different carbon sources to mitochondrial oxidative phosphorylation in cells provided with a complete medium. A shift in substrate preference towards FA at the expense of glucose distinguished late-stage hiPSC-CM from early-stage cells. A similar remodeling is observed in early postnatal heart development (Lopaschuk and Jaswal, 2010). It was also notable that glutamine made a significant contribution to substrate utilization both in early-stage and late-stage hiPSC-CM. Beyond the changes in substrate preference, we observed an overall increase in the capacity to oxidize any individual substrate, evidenced by higher mitochondrial maximal respiration and spare respiratory capacity for all substrates in late-stage hiPSC-CM. This may in part be related to an increased expression and activity of key enzymes involved in the utilization of glucose and FA. Stage-specific regulation of PDHα phosphorylation by PDKs and PDPs, which determines the entry of glucose-derived pyruvate into the TCA cycle, is well established in the perinatal heart (Sugden, et al., 2000; Sugden and Holness, 2003). We observed a reduction in inhibitory phosphorylation of PDHα in late-stage hiPSC-CM. PDK4 levels also increased 3-fold in late-stage hiPSC-CM, similar to the situation in the early postnatal heart (Chung, et al., 2010). The increase in FA oxidation in late-stage hiPSC-CM was associated with increased expression of PPARα, the master regulator of FA metabolic enzymes. In line with this, the protein levels of several downstream targets of PPARα signaling such as the FA transporter CD36 and enzymes involved in FA oxidation were increased in late-stage hiPSC-CM. Taken together, these observations indicate an integrated maturation to an early post-natal metabolic profile in late-stage hiPSC-CM.

In the developing heart, there is an increase in CM mitochondrial content as well as a structural and functional specialization, with the increase in mitochondrial content occurring mainly prenatally and in the very early postnatal phases (Pohjoismäki and Goffart, 2017; Lehman et al., 2000). In our model, the mitochondrial fraction assessed by MitoTracker Green® staining or mtDNA content was not significantly different between early-stage and late-stage hiPSC-CM. We also found no differences in the protein levels of electron transport chain components nor in the levels and post-translational phosphorylation of PCG1α, a key regulator of mitogenesis. This suggests that the major increase in mitochondrial content may have occurred before the sixth week of differentiation from iPSC (early-stage). However, we found evidence by electron microscopy of increased mitochondrial network formation in late-stage hiPSC-CM. In addition, the mitochondrial membrane potential was significantly more negative in late-stage hiPSC-CM along with a higher basal calcium concentration in the matrix. These results indicate that an important component of the increased respiratory capacity and OCR of late-stage hiPSC-CM can be ascribed to a maturation of mitochondrial structure and function.

Although the differentiation method employed in the study is optimal for the generation of 2-dimensional (2D) cultures of CM from hiPSC, the preparation remains different from the heart in some important respects. CM *in situ* in the heart are in a 3D environment and are subjected to a constant workload whereas the hiPSC-CM model is largely unloaded. The metabolic profile under loaded conditions may therefore be different. Furthermore, persistent loading of hiPSC-CM may itself alter their maturation (Kolanowski et al., 2017; Machiraju and Greenway, 2019). Future studies in hiPSC-CM-derived engineered heart muscle could address this question. Paracrine and functional interactions among different cell types in the multicellular heart may also alter the metabolic profile. Nevertheless, the 2D hiPSC-CM model is amenable to pathophysiological and pharmacological manipulations to model in vivo conditions to some extent.

In conclusion, the data presented herein provide insights into the alterations in metabolic profile of maturing hiPSC-CM. The progressive restructuring of cardiac-specific features in these cells indicates their potential to overcome the lack of full maturation previously reported. The metabolic roadmap provided here may assist in the design and interpretation of pathophysiological and pharmacological studies in hiPSC-CM where metabolism plays an important role.

## Experimental procedures

### Culture of hiPSC and CM differentiation

The hiPSC lines used in the study, FB2 (Wolf et al., 2013) and Ctrl2 (Borchert et al., 2017), were both derived from healthy donors and were described and characterized previously (Borchert et al., 2017; Wolf et al., 2013). Cells were maintained in E8 medium on Geltrex®-coated vessels and split every second day by non-enzymatic dissociation (Versene). Cultures were maintained at 37°C, 5% CO_2_. CM were differentiated using a protocol of Wnt modulation and metabolic selection as described previously (Lian et al., 2013; Tohyama et al., 2013). The differentiation protocol is illustrated schematically in Suppl. Figure 1.

### mRNA expression

Total RNA was isolated from cells using the ReliaPrep™ tissue system kit (Promega; Z6012). cDNA was produced from 1 μg of RNA sample using M-MLV reverse transcriptase (Promega; M-1302). Real time qPCR was preformed using a QuantiNova SYBR Green kit (Qiagen; 208052) on an ABI StepOne thermocycler. Mean Ct were normalized using β-2-microglobulin (*B2M*) expression. Primer sequences are reported in Suppl. Table 1.

### Immunofluorescence and flow cytometry

For immunostaining, cells were washed with PBS and fixed in formalin-free Histofix® (Roth; P087.1) for 20 minutes at room temperature. Blocking and antibody incubations were performed in 2% w/v BSA, 0.2% v/v Triton™ X-100 in PBS. 3% v/v NGS was added for antibodies raised in goat. Images were captured using an Axiovision Rel4.8-controlled Zeiss inverted fluorescence microscope.

For flow cytometry, cells were washed with PBS and incubated with 0.25% trypsin-EDTA until complete detachment. 10% FBS containing medium was added to the cells which were strained through a 22μm filter. At least 1.5×10^5^ cells per condition were used. Cells were washed with PBS, then fixed in 4% PFA for 20 minutes at room temperature. Incubations with primary antibodies were performed in 0.1% Triton X, 1% BSA in PBS, overnight at 4°C. After washing, cells were resuspended in secondary antibody solutions for 45 minutes at room temperature in the dark. After antibodies were washed off via centrifugation, samples were resuspended in PBS, and run through an Accuri C6 flow cytometer. The primary antibodies used are listed in Suppl. Table 2.

### Western blotting

Cells were lysed in RIPA buffer with protease and phosphatase inhibitors. Total protein was quantified with a Pierce™ BCA assay (Thermo Fisher). Samples were diluted to 1 μg/ml in loading buffer (20 % v/v glycerol, 4 % w/v SDS, 250 mM Trizma® Base pH 6.8, 1 mM bromophenol blue and 0.5 M 1,4-dithiothreitol). They were boiled at 95°C for 5 minutes. SDS-PAGE was performed using Bolt™ precast gels (Thermo Fisher, NW04120) and MES Buffer. Proteins were transferred onto nitrocellulose blotting membranes. Antibodies were applied overnight at 4°C on TBST-based buffers. Results were analyzed using a Licor Odyssey CLx instrument. The antibodies used are listed in Suppl. Table 3.

### Extracellular flux analysis

Studies were performed using a Seahorse xF24 analyzer. Cells were plated on Geltrex®-coated 24-well microplates at 5×10^4^ cells/well, 1 week before the assay. Cartridges were hydrated following the manufacturer’s instructions. Media were prepared on the day of each experiment, using a DMEM basal medium without glucose, glutamine or phenol red. Appropriate substrates were added as described for each assay (10 mM D-glucose, 2 mM sodium-pyruvate, 4 mM sodium-lactate) and media were brought to pH 7.4 at 37°C. Cells were equilibrated in assay medium for 45 minutes before the experiment. Three distinct assays were performed (Suppl. Table 4) and the following drugs were employed: a complex V inhibitor, oligomycin (Sigma, 75351); a mitochondrial pyruvate carrier (MPC) inhibitor, UK5099 (Sigma, PZ0160); a glutaminase inhibitor, Bis-2-(5-phenylacetamido-1,3,4-thiadiazol-2-yl)ethyl sulfide (BPTES) (Sigma, SML0601); an irreversible O-Carnitine palmitoyltransferase-1 (CPT1) inhibitor, Etomoxir (Sigma, E1905); the protonophore, Carbonyl cyanide 4-(trifluoromethoxy)phenylhydrazone (FCCP) (Sigma, C2920); a complex III inhibitor, antimycin A (Sigma, A8674); a complex I inhibitor, rotenone (Sigma, R8875). For assays involving FA oxidation, cells were starved overnight in a medium containing 2 mM L-glutamine and supplemented with 0.5 mM D-glucose and 0.5 mM L-carnitine hydrochloride. 1 mM BSA-conjugated palmitate was added to the assay medium immediately before the assay and the FCCP concentration was raised to 1 μM.

In some experiments, cells were fixed using 4% PFA for 15 minutes after completion of assays in order to perform cell-number normalization. They were then treated with 2 μg/ml DAPI for 10-15 minutes to visualize nuclei. UV-fluorescence nuclear counting was automated on an Olympus IX81 microscope and used as an index of cell number for data normalization.

### ^1^H-^13^C-nuclear magnetic resonance (NMR) to estimate metabolic flux

We performed some studies using uniformly ^13^C-labeled glucose (^13^C-U glucose) and NMR to estimate the activity of different glucose-utilizing pathways in hiPSC-CM (Schnelle et al., 2020). Cells (>5×10^6^ cells per sample) were incubated with ^13^C-U glucose (10 mM) for 16 hours. Cells were kept on dry ice and scraped in methanol. Chloroform and water were added and the polar phase was isolated after cold centrifugation. Samples were dried and subsequently reconstituted in P-buffer (40 mM NaH_2_PO_4_, 60 mM Na_2_HPO_4_, 0.01 % w/v DSS, 1.5 mM Imidazole, in D_2_O). They were analyzed at 25 ^o^C on a Bruker Avance 700 MHz NMR spectrometer. The ^13^C labeled metabolites were calculated using heteronuclear single quantum coherence (HSQC) as described previously (Schnelle et al., 2020). Spectra were processed using the MetaboLab software package (Ludwig and Günther, 2011).

### Transmission electron microscopy

Cells grown on Thermonox coverslips were fixed in 2% glutaraldehyde in 0.1 M sodium cacodylate buffer, pH 7-7.4, for 60 minutes (room temperature) followed by a post-fixation step with 1% osmium tetroxide in 0.1 M sodium cacodylate buffer for 1 hour (room temperature). Following fixation the cells were dehydrated in a graded series of ethanol, equilibrated with propylene oxide, and infiltrated with epoxy resin (Agar100 from Agar Scientific) using gelatin molds and cured at 60°C for 48 hours. Coverslips were removed from the resin blocks using liquid nitrogen and 70-80 nm ultrathin sections were prepared with an Ultracut E ultramicrotome (Reichert-Jung, Leica Microsystems Ltd, Milton Keynes, UK), mounted on copper grids, and contrasted with uranyl acetate and lead citrate. Samples were examined on a JOEL JEM 1400Plus Transmission Electron Microscope operated at 120 kV, and images acquired with a high sensitivity sCMOS camera at the JEOL Centre for Ultrastructural Imaging (CUI).

### Mitochondrial parameters

To calculate the mitochondrial fraction in live cells, they were loaded with MitoTracker® Green (1:1000 in culture medium for 30 minutes). Cells were then imaged by epifluorescence and the mitochondrial fraction reported as signal area/total cell area.

Mitochondrial DNA (mtDNA) content was normalized to nuclear DNA (nDNA). nDNA was obtained via qPCR on genomic DNA isolated from cells harvested in 100 mM Tris-HCl, pH 8; 5 mM EDTA, pH 8; 0.2 % w/v SDS; 200 nM NaCl. *B2M* was used as nuclear DNA reference. Primers for mtUUR were: forward, CACCCAAGAACAGGGTTTGT; reverse, TGGCCATGGGTATGTTGTTA.

Mitochondrial membrane potential was assessed using FACS in cells loaded with tetramethylrhodamine ethyl ester (TMRE) for 30 minutes in culture medium (Beretta et al., 2020). Mitochondrial calcium levels were estimated using cells transfected with a mitochondrial-targeted Cameleon FRET probe as previously described (Beretta et al., 2020). FRET measurements were performed on a Nikon A1R confocal microscope.

### Statistical analysis

Data are expressed as mean ± S.E.M. Statistical analyses were performed using GraphPad Prism 7. Groups were compared by 1-way or 2-way analyses of variance (ANOVA) followed by Tukey’s multiple comparisons tests, or unpaired twotailed Student’s t-tests as appropriate. p<0.05 was considered as the threshold for statistical significance.

## Acknowledgements

This work was supported by the British Heart Foundation (RE/18/2/34213 and CH/1999001/11735; AMS); a Fondation Leducq Transatlantic Network of Excellence award (17CVD04; AMS); the Department of Health via a National Institute for Health Research (NIHR) Biomedical Research Centre award to Guy’s & St Thomas’ NHS Foundation Trust in partnership with King’s College London (IS-BRC-1215-20006; AMS); the Deutsche Forschungsgemeinschaft through the International Research Training Group Award (IRTG) 1816 (KSB, AMS); and the German Center for Cardiovascular Research (DZHK; KSB). GE was a Joint PhD student under IRTG 1816. We thank Roland Fleck and the Centre of Ultrastructural Imaging at King’s College London for access to electron microscopy.

## Author contributions

GE: conceptualization, data curation, formal analysis, Investigation, methodology, visualization, writing - original draft; AZ: conceptualization, Investigation, supervision; AP: formal analysis, investigation, visualization; MC: formal analysis, investigation, validation; MB: investigation, resources; KB: investigation, visualization, validation; KSB: conceptualization, funding acquisition, methodology, supervision, validation; AMS: conceptualization, funding acquisition, overall project supervision, writing – review & editing.

## Supplementary experimental procedures

**Supplementary Table 1.**
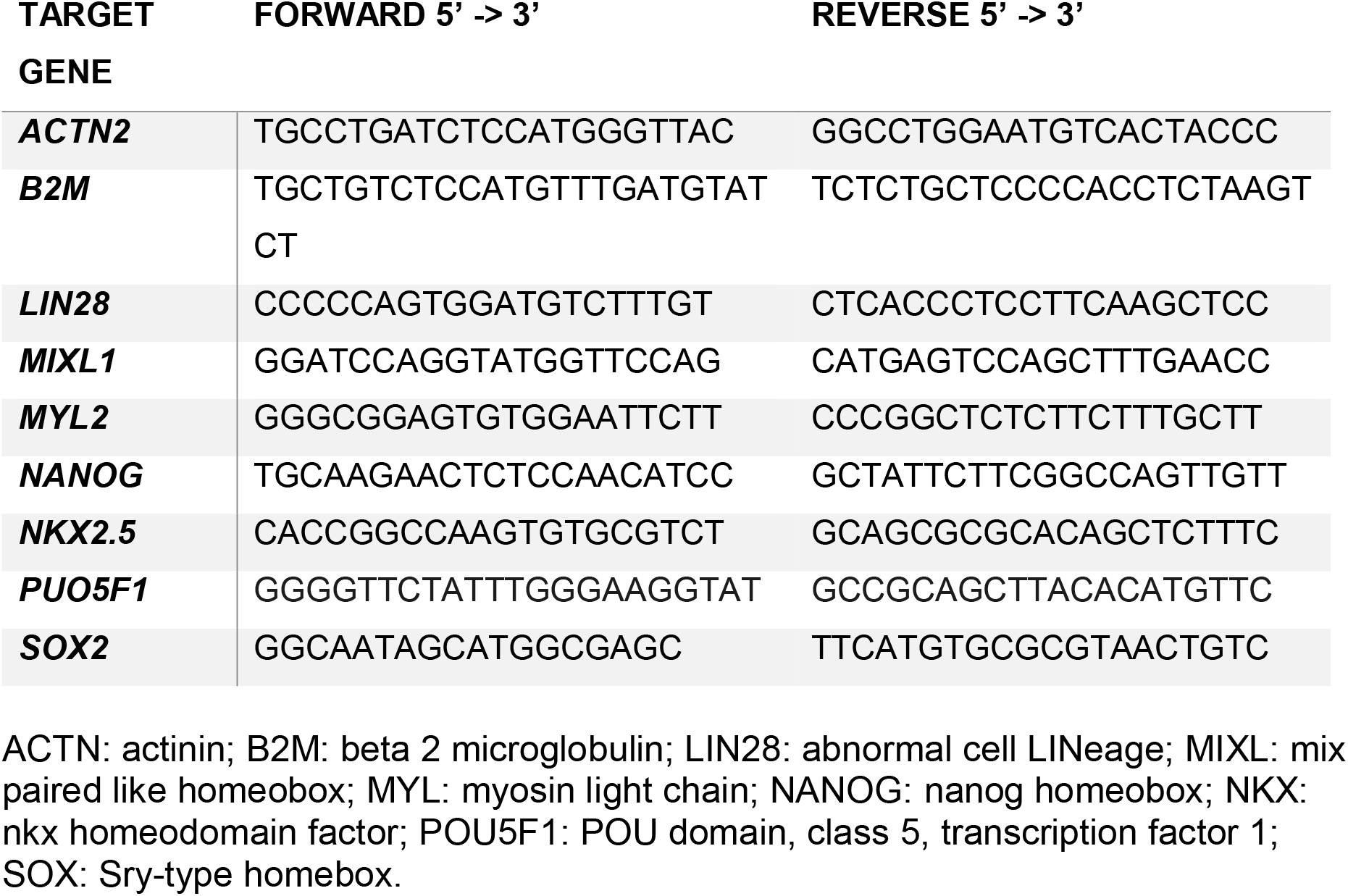
Primer sequences for RT-qPCR

**Supplementary Table 2.** Antibodies for immunostaining and FACS

| **TARGET**<br>**PROTEIN** | **SPECIES** | **DILUTION** | **CATALOGUE #** |
| --- | --- | --- | --- |
| **ACTININ** | Rabbit | 1:500<br>(1:750 FACS) | Abcam, ab68194 |
| **cTNT** | Mouse | 1:750 | Thermo Fisher, MS-295-PAbx |
| **LIN28** | Goat | 1:300 | R&D, AF3757 |
| **MLC2A** | Mouse | 1:333<br>(1:250 FACS) | Abcam, ab68086 |
| **MLC2V** | Rabbit | 1:333<br>(1:250 FACS) | Proteintech, 10906-1-AP |
| **NANOG** | Rabbit | 1:100 | Thermo Fisher, PA1-097 |
| **OCT4** | Goat | 1:500 | R&D, AF1759 |
| **SSEA-4** | Mouse | 1:200 | Abcam, ab16287 |
| **SOX2** | Mouse | 1:200 | R&D, MAB2018 |
| **TITIN C-TERM** | Rabbit | 1:750 | Myomedix, TTN-M |
| **TITIN N-TERM** | Rabbit | 1:750 | Myomedix, TTN-Z |
| **TRA-1-60** | Mouse | 1:200 | Abcam, ab16288 |

cTNT: cardiac troponin T; LIN: abnormal cell LiNeage; MLC: myosin light chain; OCT: octamer binding transcription factor; SSEA: stage specific embryo antigen; SOX: Sry-type homeobox; TRA: T cell receptor alpha locus.

**Supplementary Table 3.**
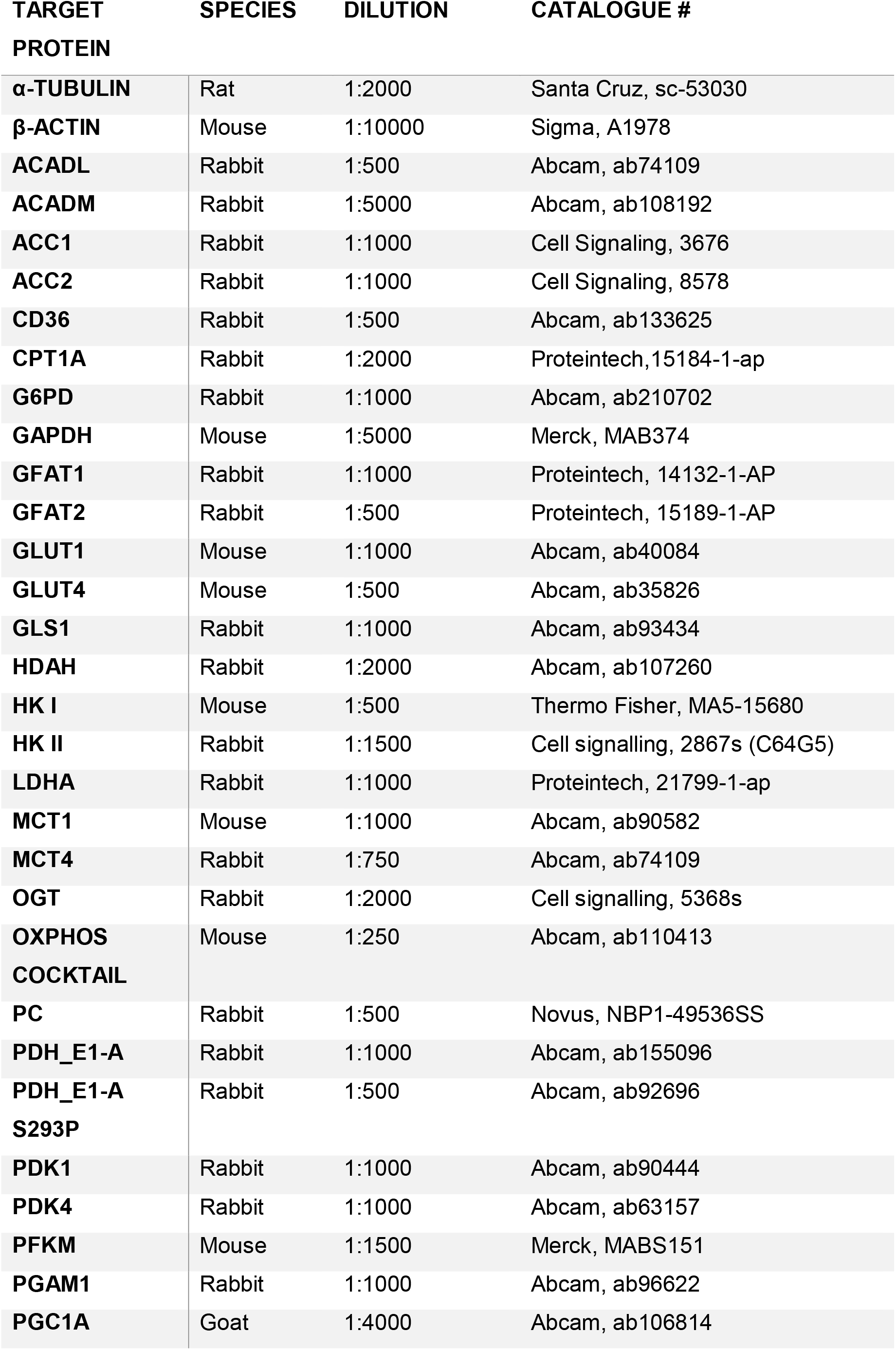
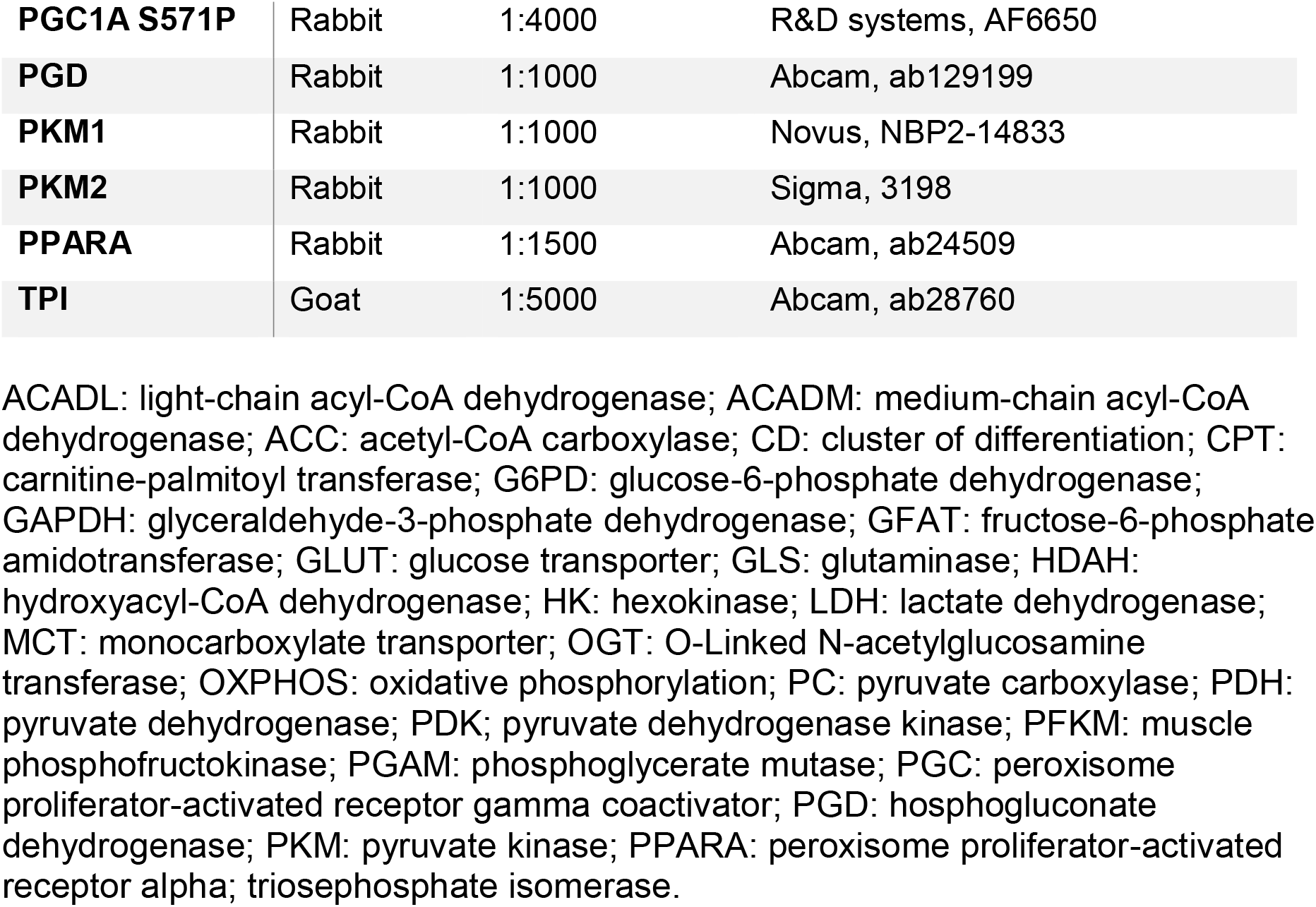
Antibodies for western blotting

| **TARGET**<br>**PROTEIN** | **SPECIES** | **DILUTION** | **CATALOGUE #** |
| --- | --- | --- | --- |
| **α-TUBULIN** | Rat | 1:2000 | Santa Cruz, sc-53030 |
| **β-ACTIN** | Mouse | 1:10000 | Sigma, A1978 |
| **ACADL** | Rabbit | 1:500 | Abcam, ab74109 |
| **ACADM** | Rabbit | 1:5000 | Abcam, ab108192 |
| **ACC1** | Rabbit | 1:1000 | Cell Signaling, 3676 |
| **ACC2** | Rabbit | 1:1000 | Cell Signaling, 8578 |
| **CD36** | Rabbit | 1:500 | Abcam, ab133625 |
| **CPT1A** | Rabbit | 1:2000 | Proteintech, 15184-1-ap |
| **G6PD** | Rabbit | 1:1000 | Abcam, ab210702 |
| **GAPDH** | Mouse | 1:5000 | Merck, MAB374 |
| **GFAT1** | Rabbit | 1:1000 | Proteintech, 14132-1-AP |
| **GFAT2** | Rabbit | 1:500 | Proteintech, 15189-1-AP |
| **GLUT1** | Mouse | 1:1000 | Abcam, ab40084 |
| **GLUT4** | Mouse | 1:500 | Abcam, ab35826 |
| **GLS1** | Rabbit | 1:1000 | Abcam, ab93434 |
| **HDAH** | Rabbit | 1:2000 | Abcam, ab107260 |
| **HK I** | Mouse | 1:500 | Thermo Fisher, MA5-15680 |
| **HK II** | Rabbit | 1:1500 | Cell signalling, 2867s (C64G5) |
| **LDHA** | Rabbit | 1:1000 | Proteintech, 21799-1-ap |
| **MCT1** | Mouse | 1:1000 | Abcam, ab90582 |
| **MCT4** | Rabbit | 1:750 | Abcam, ab74109 |
| **OGT** | Rabbit | 1:2000 | Cell signalling, 5368s |
| **OXPHOS**<br>**COCKTAIL** | Mouse | 1:250 | Abcam, ab110413 |
| **PC** | Rabbit | 1:500 | Novus, NBP1-49536SS |
| **PDH_E1-A** | Rabbit | 1:1000 | Abcam, ab155096 |
| **PDH_E1-A**<br>**S293P** | Rabbit | 1:500 | Abcam, ab92696 |
| **PDK1** | Rabbit | 1:1000 | Abcam, ab90444 |
| **PDK4** | Rabbit | 1:1000 | Abcam, ab63157 |
| **PFKM** | Mouse | 1:1500 | Merck, MABS151 |
| **PGAM1** | Rabbit | 1:1000 | Abcam, ab96622 |
| **PGC1A** | Goat | 1:4000 | Abcam, ab106814 |
| **PGC1A S571P** | Rabbit | 1:4000 | R&D systems, AF6650 |
| **PGD** | Rabbit | 1:1000 | Abcam, ab129199 |
| **PKM1** | Rabbit | 1:1000 | Novus, NBP2-14833 |
| **PKM2** | Rabbit | 1:1000 | Sigma, 3198 |
| **PPARA** | Rabbit | 1:1500 | Abcam, ab24509 |
| **TPI** | Goat | 1:5000 | Abcam, ab28760 |

ACADL: light-chain acyl-CoA dehydrogenase; ACADM: medium-chain acyl-CoA dehydrogenase; ACC: acetyl-CoA carboxylase; CD: cluster of differentiation; CPT: carnitine-palmitoyl transferase; G6PD: glucose-6-phosphate dehydrogenase; GAPDH: glyceraldehyde-3-phosphate dehydrogenase; GFAT: fructose-6-phosphate amidotransferase; GLUT: glucose transporter; GLS: glutaminase; HDAH: hydroxyacyl-CoA dehydrogenase; HK: hexokinase; LDH: lactate dehydrogenase; MCT: monocarboxylate transporter; OGT: O-Linked N-acetylglucosamine transferase; OXPHOS: oxidative phosphorylation; PC: pyruvate carboxylase; PDH: pyruvate dehydrogenase; PDK: pyruvate dehydrogenase kinase; PFKM: muscle phosphofructokinase; PGAM: phosphoglycerate mutase; PGC: peroxisome proliferator-activated receptor gamma coactivator; PGD: phosphogluconate dehydrogenase; PKM: pyruvate kinase; PPARA: peroxisome proliferator-activated receptor alpha; triosephosphate isomerase.

**Supplementary Table 4.** Summary of protocols for Seahorse analyzer

|  | Glycolysis stress test |  | Mitochondrial Stress test |  | Fuel Source test |  |
| --- | --- | --- | --- | --- | --- | --- |
| Medium | Substrate free |  | Single-substrate |  | Complete |  |
| Injection 1 | Glucose | 10 mM | Oligomycin A | 1 μM | UK5099 or BPTES or Etomoxir | 7.5 μM<br>10 μM<br>15 μM |
| Injection 2 | Oligomycin A | 1 μM | FCCP | 0.5 μM | Combination | |
| Injection 3 | 2-Deoxy-D-Glucose | 100 mM | Antimycin A + Rotenone | 1 μM<br>1 μM | Antimycin A + Rotenone | 1 μM<br>1 μM |

**Supplementary Figure 1.**
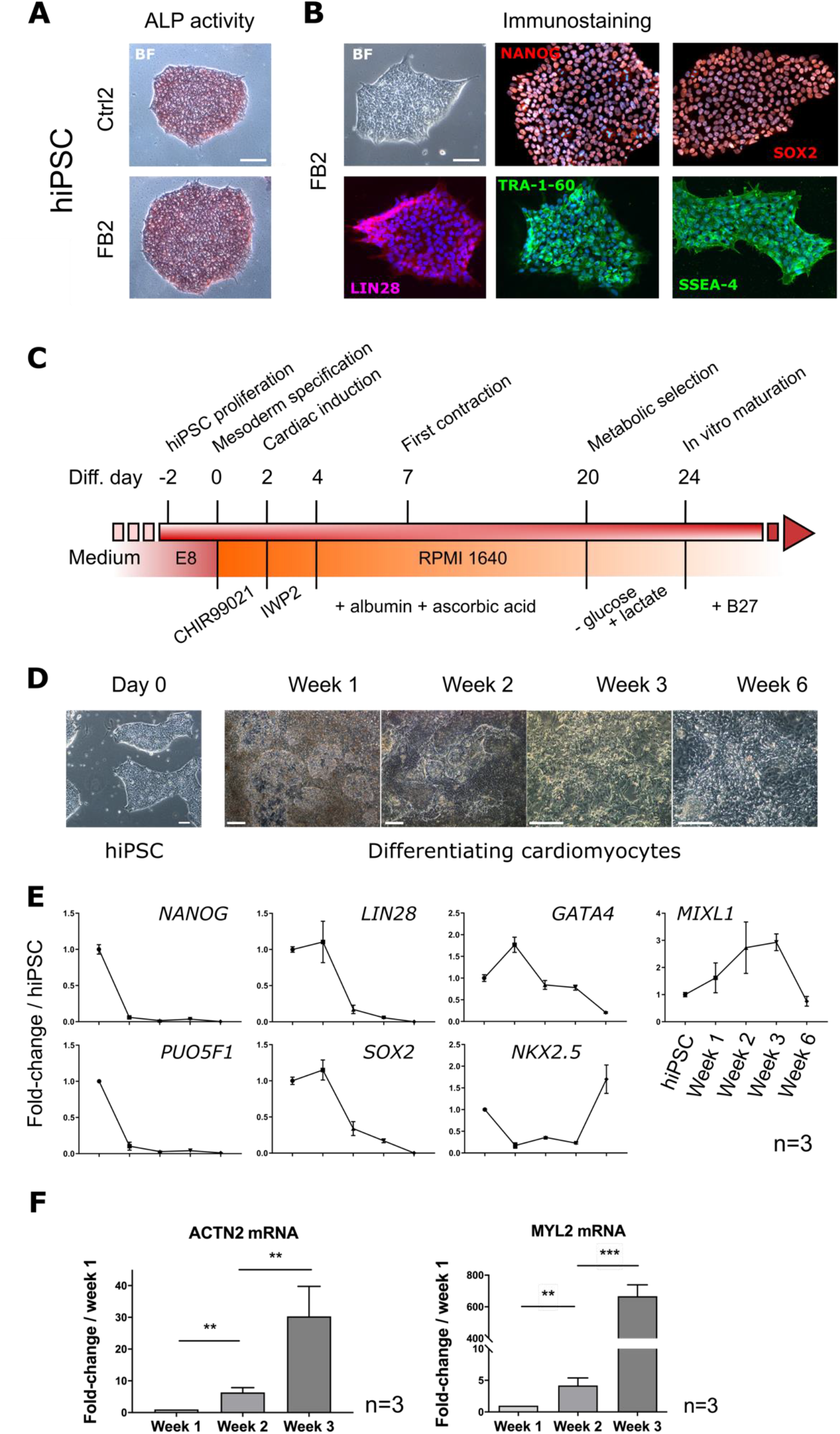
Characterization of hiPSC differentiation to cardiomyocytes. A) Representative images of alkaline phosphatase (ALP) activity in the two hiPSC lines utilized in this study (FB2 and Ctrl2). Red staining, positive for ALP activity. Scale bar, 100 μm. B) Representative images of immunofluorescence for markers of pluripotency (NANOG, SOX2, LIN28, TRA-1-60, SSEA4). Scale bar, 200 μm. BF: bright field. C) Schematic of the hiPSC-CM differentiation protocol. CHIR99021: small molecule inhibitor of glycogen-synthase kinase subunit 3; IWP2: small molecule inhibitor of Wnt signaling by inactivating the Porcupine O-acyltransferase which palmitoylates Wnt proteins; Albumin: human recombinant; B27: commercially available growth supplement. D) Brightfield images of the timecourse of hiPSC-CM differentiation. Scale bar, 200 μm. E) Time-course of changes in mRNA levels of pluripotency (*NANOG, LIN28, POU5F1, SOX2*) and cardiac-specification markers (*GATA4, NKX2.5, MIXL1*) in differentiating hiPSC-CM. Data are mean ± S.E.M. Significances by 1-way ANOVA and Tukey’s test for multiple comparisons are shown in Suppl. Table 5; n=3 differentiation experiments F) Timecourse of changes in mRNA levels of cardiac sarcomeric proteins in differentiating hiPSC-CM. Data are mean ± S.E.M. **, p<0.01, ***, p<0.001 by 1-way ANOVA and Tukey’s test; n=3 differentiation experiments.

**Supplementary Figure 2.**
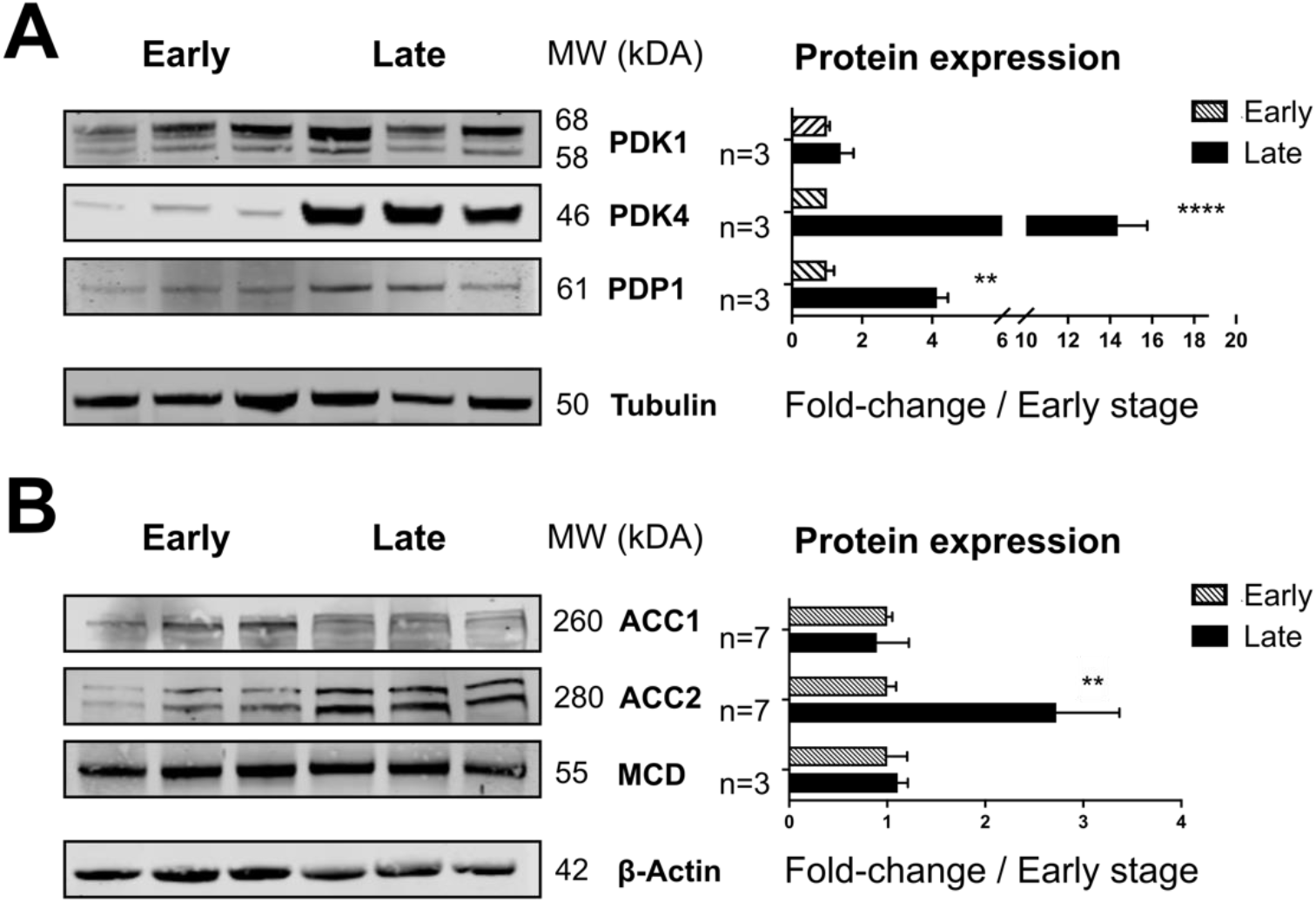
Changes in protein levels of enzymes involved in flux into the TCA. A-B) Left panels are representative western blots for the proteins shown. Right panels show quantification of relative protein expression as foldchange compared to early-stage. Data are mean ± S.E.M. *, p<0.05; **, p<0.01; ****, p<0.0001 by unpaired t-test. 3 ⩽ n ⩽ 7 as indicated in figure. PDK, pyruvate dehydrogenase kinase; ACC, acetyl-CoA carboxylase; MCD, malonyl-CoA decarboxylase.

